# Control of microglial dynamics by Arp2/3 and the autism and schizophrenia-associated protein Cyfip1

**DOI:** 10.1101/2020.05.31.124941

**Authors:** James Drew, I. Lorena Arancibia-Carcamo, Renaud Jolivet, Guillermo Lopez-Domenech, David Attwell, Josef T. Kittler

## Abstract

Microglia use a highly complex and dynamic network of processes to sense and respond to their surroundings. Microglial dynamics differ throughout development and in neurological and neuropsychiatric disease, though mechanistic insight into these changes is lacking. Here we identify novel roles for regulators of the actin cytoskeleton in controlling microglial behaviour. We show that the actin branching complex Arp2/3 is critical for maintaining microglial morphology and required for surveillance but not chemotactic motility. Neuropsychiatric disease-associated Cyfip1, a core component of the WAVE regulatory complex that links Rac1 signalling to Arp2/3 activation, is highly expressed in microglia but has unknown function. We report that conditional deletion of Cyfip1 in mouse microglia reduces morphological complexity and surveillance of brain parenchyma, and increases activation state as defined by CD68 expression. Thus, altered actin-dependent microglial dynamics mediated by Cyfip1 and Arp2/3 may contribute to neuropsychiatric disease.

## Introduction

Microglia are the resident immune cells of the brain, performing innate immune functions during infection or damage. Imaging studies have shown that ‘resting’ microglia in the healthy brain are extremely motile, constantly surveying their surroundings using a complex network of branched processes (Davalos et al., 2005; Nimmerjahn et al., 2005; Wake et al., 2009). Subsequent research has revealed roles for microglia in shaping many aspects of brain development, including neuron survival, proliferation, migration, and synapse formation and pruning (Lehrman et al., 2018; Miyamoto et al., 2016; Neniskyte and Gross, 2017; Parkhurst et al., 2013; Thion et al., 2018). In parallel, clinical evidence suggests a pathological role for microglia in neurodegenerative and neuropsychiatric disorders, in which neuronal connectivity is often disrupted (Estes and McAllister, 2015; Sekar et al., 2016; Sellgren et al., 2019; Velmeshev et al., 2019). How microglia generate ramified, dynamic protrusions and how this relates to the pathology of neurological disorders remains poorly understood.

The actin cytoskeleton is required for cells to move and generate complex morphologies. Cells use branched actin networks to form lamellipodia, sheet-like protrusions that distort the plasma membrane during migration and growth (Krause and Gautreau, 2014; Ridley, 2011; Rougerie et al., 2013; Sossey-Alaoui et al., 2007). These branched networks are produced by the actin-related complex Arp2/3, which generates new filamentous actin branches from existing F-actin (Krause and Gautreau, 2014; Mullins et al., 1997). The Arp2/3 complex is basally inactive but can be switched on by interaction with the WAVE regulatory complex (WRC), which binds directly to Arp2/3 to promote actin branching, membrane protrusion and cell migration (Ridley, 2011). The WRC is a heteropentameric complex composed of paralogs of the proteins Cyfip, Abi, Nckap1, Wave and Hspc300. Interestingly, rare variants and truncations of WRC components *WAVE1*, *NCKAP1*, *CYFIP1* and *CYFIP2* are genetically linked to autistic spectrum disorders (ASDs), epilepsy and intellectual disability, while *ABI3* and *CYFIP2* are associated with Alzheimer’s disease - implicating regulation of actin dynamics by the WRC in susceptibility to neurological disease (Sims et al., 2017; Wang et al., 2016; Zweier et al., 2019; Tiwari et al., 2016).

Cytoplasmic FMRP-interacting protein 1 (Cyfip1) is one of two Cyfip proteins in mammalian systems, and couples activation of the WRC and Arp2/3 to upstream signaling from the small RhoGTPase Rac1 (Chen et al., 2017, 2010; Koronakis et al., 2011). Human *CYFIP1* is of clinical interest due to its strong associations with neuropsychiatric disease. The gene resides in the 15q11.2 region of the genome that is vulnerable to interstitial copy number variations (CNVs), the smallest of which comprises *NIPA1*, *NIPA2*, *TUBGCP5* and *CYFIP1* (Cox and Butler, 2015). Both deletions and duplications of 15q11.2 have been linked to schizophrenia, ASDs, intellectual disability, epilepsy, ADHD and neurodevelopmental delay, and are correlated with changes in *CYFIP1* expression (Kirov et al., 2012; Marshall et al., 2017; Stefansson et al., 2008, 2014; van der Zwaag et al., 2010). Associations between schizophrenia and variant mutations in *CYFIP1* have also been reported (Waltes et al., 2014; Wang et al., 2015; Yoon et al., 2014; Zhao et al., 2013). Cyfip1 deletion or haploinsufficiency in mice leads to neuronal morphology, plasticity and connectivity defects (Cioni et al., 2018; Davenport et al., 2019; Domínguez-Iturza et al., 2019; Pathania et al., 2014; De Rubeis et al., 2013; Hsiao et al., 2016). Despite a focus on neuronal functions of Cyfip1, transcriptomic analyses of both human and murine tissues have shown that the gene is widely expressed in glia, particularly microglia (Zeisel et al., 2018; Zhang et al., 2014, 2016). Given their emerging role in the pathology of neuropsychiatric disease, whether Cyfip1 has important cell-autonomous functions in glial cells that could contribute to its clinical associations is a pressing question (Vainchtein and Molofsky, 2020).

Here we show that maintenance of microglial morphology is critically dependent on sustained Arp2/3 activity. Inhibition of Arp2/3 leads to a selective impairment in surveillance whilst ATP/ADP-driven chemotaxis remains intact. Using an inducible conditional Cyfip1 knockout mouse, we show that deletion of microglial Cyfip1 reduces morphological complexity, impairs surveillance motility and leads to increased microglial activation. These findings uncover a novel role for Cyfip1 and Arp2/3-associated proteins in regulating microglial form and function, and open up a new mechanistic basis for clinical associations of *CYFIP1* with neurological and neuropsychiatric disease.

## Results

### Arp2/3 activity is required to maintain ramification of microglia

It has previously been reported that actin polymerisation is required for microglial motility (Bernier et al., 2019; Hines et al., 2009). However, little is known about the intracellular signalling pathways governing actin remodelling during microglial surveillance or chemotaxis to damaged cells - two stereotyped forms of motility exhibited by microglia *in situ,* which are known to be controlled by different signalling mechanisms (Madry et al., 2018b). Interestingly, Gene Ontology pathway analysis of genes expressed more in microglia than in other brain cells reveals actin-binding genes to be overrepresented (Fig.1A-B, Supp. Table 1). To investigate this, acute hippocampal slices from mice expressing eGFP under the microglia-specific Iba1 promoter were live imaged using 2-photon microscopy (Hirasawa et al., 2005). Microglial motility during pharmacological inhibition of key actin-related targets was assessed based on published protocols (Madry et al., 2018a, 2018b). Briefly, random surveillance of the parenchyma by microglial processes was measured as the number of image pixels newly surveyed or retracted from in the 30 sec between image frames (Madry et al., 2018a), termed surveillance index, and chemotactic response was quantified as the rate of process convergence towards a laser-induced lesion (see Methods). The importance of actin dynamics was confirmed using cytochalasin D (10 μM), an inhibitor of F-actin polymerisation, which completely abolished microglial motility - preventing both surveillance (Fig.S1A-B, p<0.0001) and chemotaxis (Fig. S1E-F, p<0.001). In contrast, depolymerisation of microtubules with vinblastine (10 μM) did not affect microglial surveillance over this timeframe (Fig. S1C-D).

**Fig. 1:**
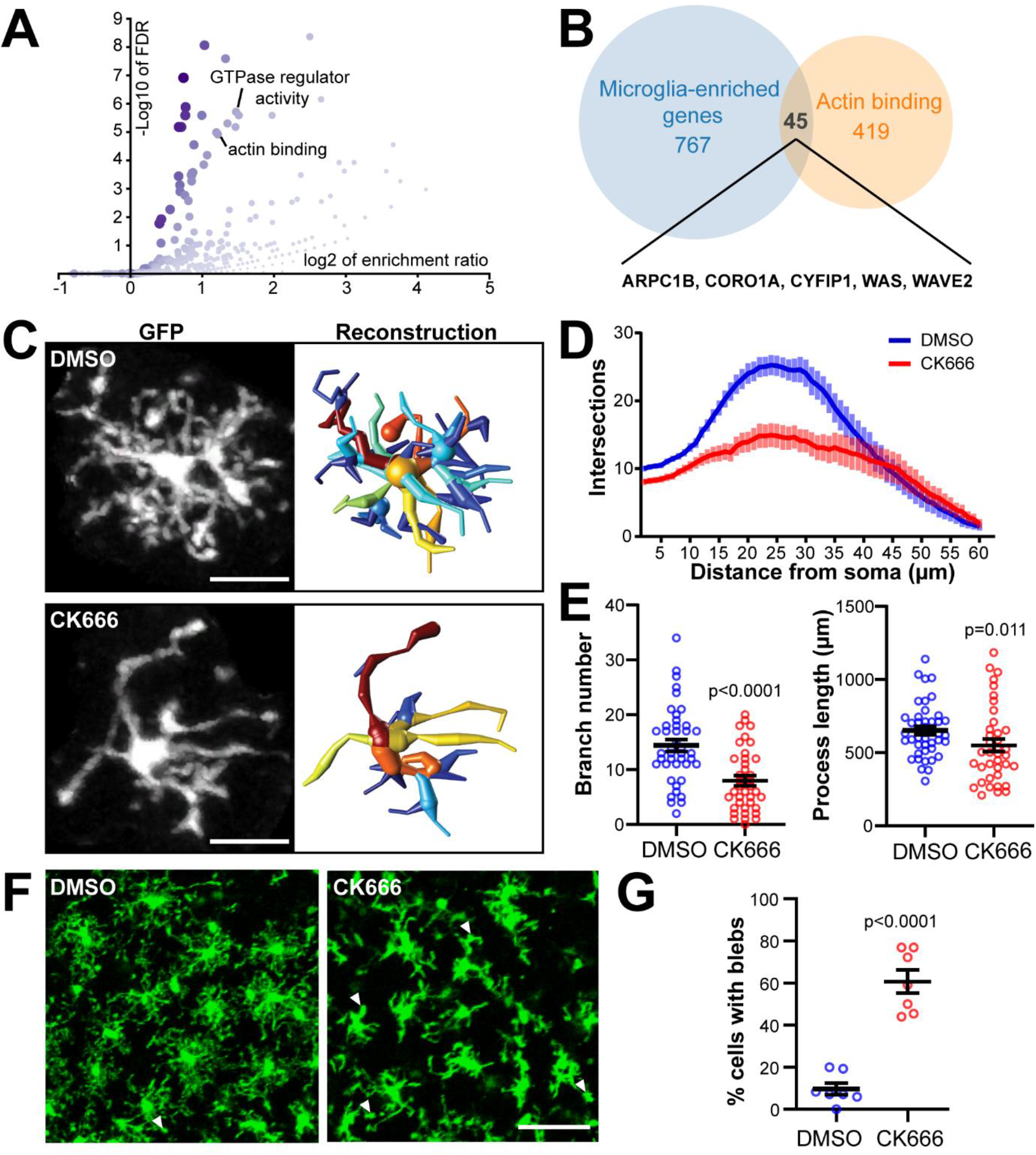
Arp2/3 activity is required to maintain ramification of microglia. **(A)** Volcano plot of GO pathways over-represented in microglia (see Supp Table 1). **(B)** 45 genes present in the ‘actin binding’ GO pathway are enriched in microglia, including Arp2/3 complex (ARPC1B) and members of the WAVE regulatory complex (CYFIP1, WASF2). **(C)** Representative cells from slices treated with DMSO (top) or 200 μM CK666 (bottom), showing loss of higher order processors. Left panel: Maximum intensity projection of GFP fluorescence. Right panel: 3D reconstruction of cell morphology with colour mapping to distinguish separate processes. Scale bar: 10 μm. **(D)** Sholl analysis of intersections from 3D reconstructions (p<0.001, 2-way repeated measures ANOVA, n=38-42 cells). **(E)** Summary of morphological characteristics; decrease in branching (DMSO: 14.4 ± 1.1, CK666: 7.9 ± 0.9, p<0.0001, Mann-Whitney) and total process length (DMSO: 651.4 ± 28.7 μm, CK666: 550.2 ± 42.7 μm, p=0.011, Mann-Whitney). **(F)** CK666-treated microglia often had enlarged cellular protrusions. Left: Field of view of CK666-treated slice highlighting ‘blebs’ (white arrows). Scale bar: 25 μm **(G)** Quantification of proportion of microglia with blebs (DMSO: 9.7 ± 2.8 %, CK666: 60.7 ± 5.5 %, p<0.0001, Student’s t-test).

Interestingly, several actin-associated genes found enriched in microglia pointed to potential roles for Arp2/3 and the upstream activators Rac1, Wasf2 and the psychiatric disease-associated WRC component Cyfip1 (Fig.1B). To investigate the effects of inhibiting Arp2/3-dependent actin branching on microglial morphology, we treated acute slices with the Arp2/3 inhibitor CK666 for 30 minutes prior to and during imaging (Hetrick et al., 2013). Inhibition of Arp2/3 led to a dramatic reduction in the complexity of microglial branching, characterised by retraction of secondary and tertiary process branches (Fig.1C). Sholl analysis of 3D reconstructions from DMSO and CK666-treated microglia after 30 minutes treatment showed a dramatic loss of process complexity (Fig.1D, p<0.001). The number of branches and total process length in CK666-treated cells was reduced (Fig.1E, branches: p<0.0001, total process length: p<0.01). Primary processes often appeared swollen, with aberrant membrane blebbing observed at process tips (Fig.1F). When quantified, approximately 60% of CK666-treated cells showed such blebbing, a 6-fold increase over control (Fig.1G, p<0.0001). Thus, Arp2/3 activity is critical for maintaining the normal ramified morphology of microglia.

### Inhibition of Arp2/3 and Rac1 leads to defects in surveillance of brain parenchyma

We hypothesised that these changes in morphology would lead to defective motility of CK666-treated microglia. Analysis of surveillance revealed an over 50% reduction in pixels surveyed by Arp2/3 inhibited cells compared to vehicle (DMSO) treatment for which surveillance remained constant over the imaging period (Fig. 2A-B, p<0.0001). Surprisingly, the chemotactic response to a laser-induced lesion remained robust under Arp2/3 inhibition (Fig.2C-E). The overall convergence of processes towards the lesion site was unchanged (Fig.2D), with no difference in the speed of individual processes observed (Fig. 2E). These data suggest that Arp2/3 activity is required for microglia to effectively survey their surroundings but is dispensable for chemotaxis.

**Fig. 2:**
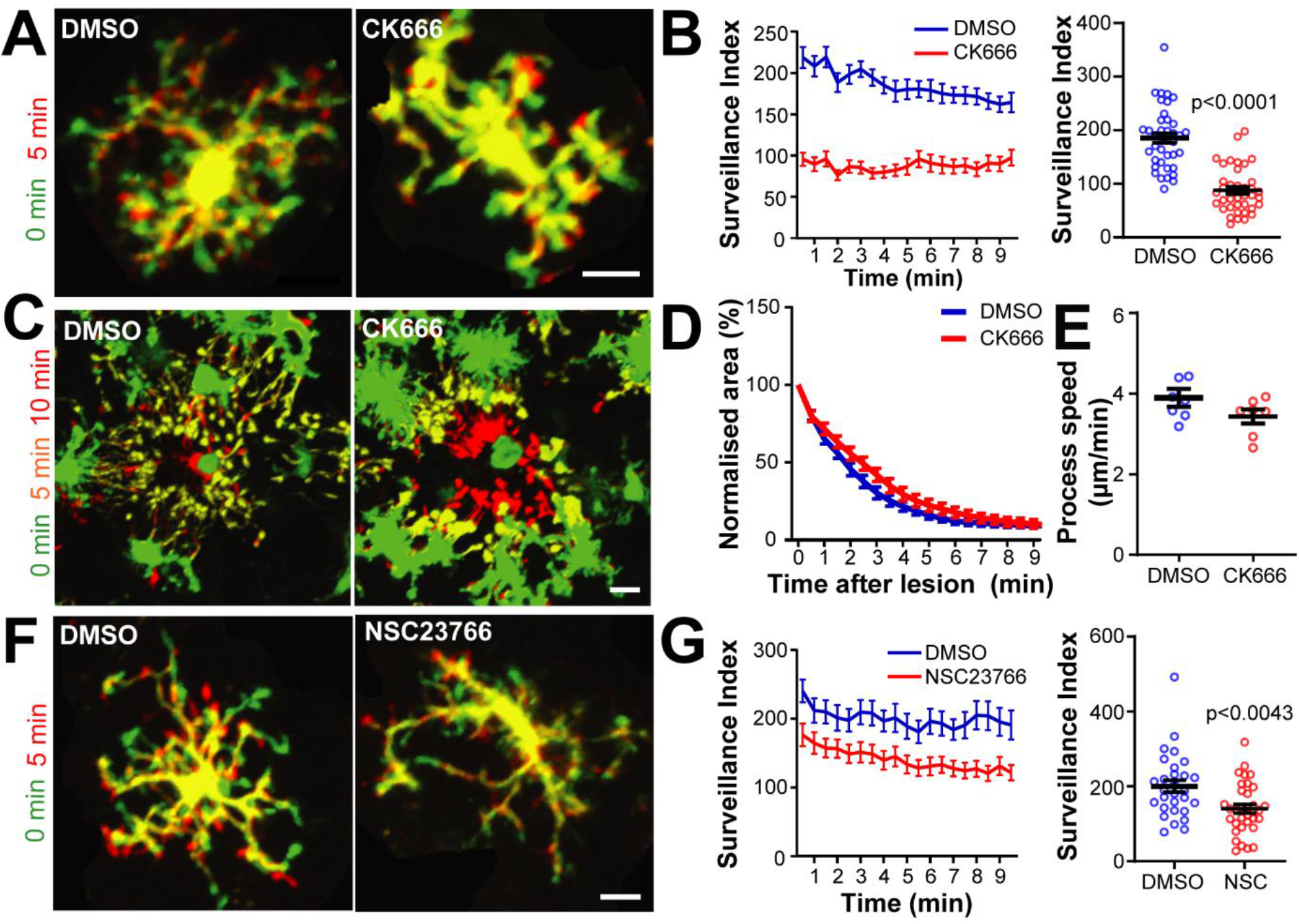
Inhibition of Arp2/3 and Rac1 leads to defects in surveillance of brain parenchyma. **(A)** Superimposed timelapse images of representative cells from DMSO or CK666 treated slices, showing extensions (red), retractions (green) and stable regions (yellow) over 5 minutes. Scale bar: 20 μm. **(B)** Quantification of surveillance from 10 minute long, 30 second frame movies. Surveillance index of CK666-treated cells decreased compared to controls (DMSO: 185.3 ± 9.2 px per 30 sec, CK666: 87.8 ± 6.9 px per 30 sec, p<0.0001, Mann-Whitney, n=38-39 cells). **(C-E)** Chemotaxis during Arp2/3 inhibition. **(C)** Superimposed timelapse showing process chemotaxis towards circular lesion in both DMSO and CK666 treated slices. Scale bar: 10 μm. **(D)** Quantification of process convergence (n=11 lesions). **(E)** Single process tracking of CK666-treated processes (DMSO: 3.9 ± 0.1 μm/min^−1^, CK666: 3.5 ± 0.1 μm/min^−1^, n= mean process speed from 7 lesions). **(F)** Superimposed timelapse images of microglia from DMSO or NSC23766 treated slices. Scale bar: 10 μm. **(G)** Surveillance index of NSC23766-treated cells decreased compared to control (DMSO: 200.5 ± 16.1 px per 30 sec, NSC23766: 140.5 ± 11.7 px per 30 sec, p<0.01, Mann-Whitney, n=29-34 cells).

Arp2/3 activity is regulated by numerous upstream pathways, one of which is Rac1-mediated activation of the WAVE regulatory complex (WRC). To investigate whether Rac1 activity was controlling microglial motility, acute hippocampal slices were pretreated with 100 μM of the Rac1 blocker NSC23766 (Hou et al., 2014). Analysis of surveillance revealed a 30% reduction in surveillance compared to DMSO-treated control slices (Fig.2 F-G). Surprisingly, the morphological complexity of the microglia appeared normal (Fig.S2A-B), with branching and process length unchanged (Fig. S2C). Bleb-like membrane protrusions were also observed in NSC23766-treated microglia (Fig. S2D-E). Thus, Rac1 inhibition recapitulates some but not all of the phenotypes observed with Arp2/3 inhibition, suggesting additional regulation of the complex by other upstream pathways.

### Generation of a microglia-specific Cyfip1 conditional knockout mouse model

Cyfip1 is central to the transmission of active Rac1 signalling to the Arp2/3 complex, via its role in the WAVE regulatory complex. Given the effects of disrupting the Rac1-Arp2/3 axis and Cyfip1’s enrichment in microglia, we hypothesised that Cyfip1 loss could alter microglial dynamic behaviours. Using published RNA sequencing from the Barres lab (Zhang et al., 2014), an analysis of microglial expression of paralogs of the different WRC components compared to other brain cells revealed a canonical ‘microglial WRC’ composed of Cyfip1, Abi3, Nap1l and Wave2 (Fig. 3A). Notably, Cyfip2 expression is virtually absent in microglia, in both humans and mice (Fig. S3), supporting the case for Cyfip1 being a critical component of the microglial WRC. Thus, we hypothesised that loss of microglial Cyfip1 could affect actin dynamics via the Rac1-WRC-Arp2/3 signalling axis.

**Fig.3:**
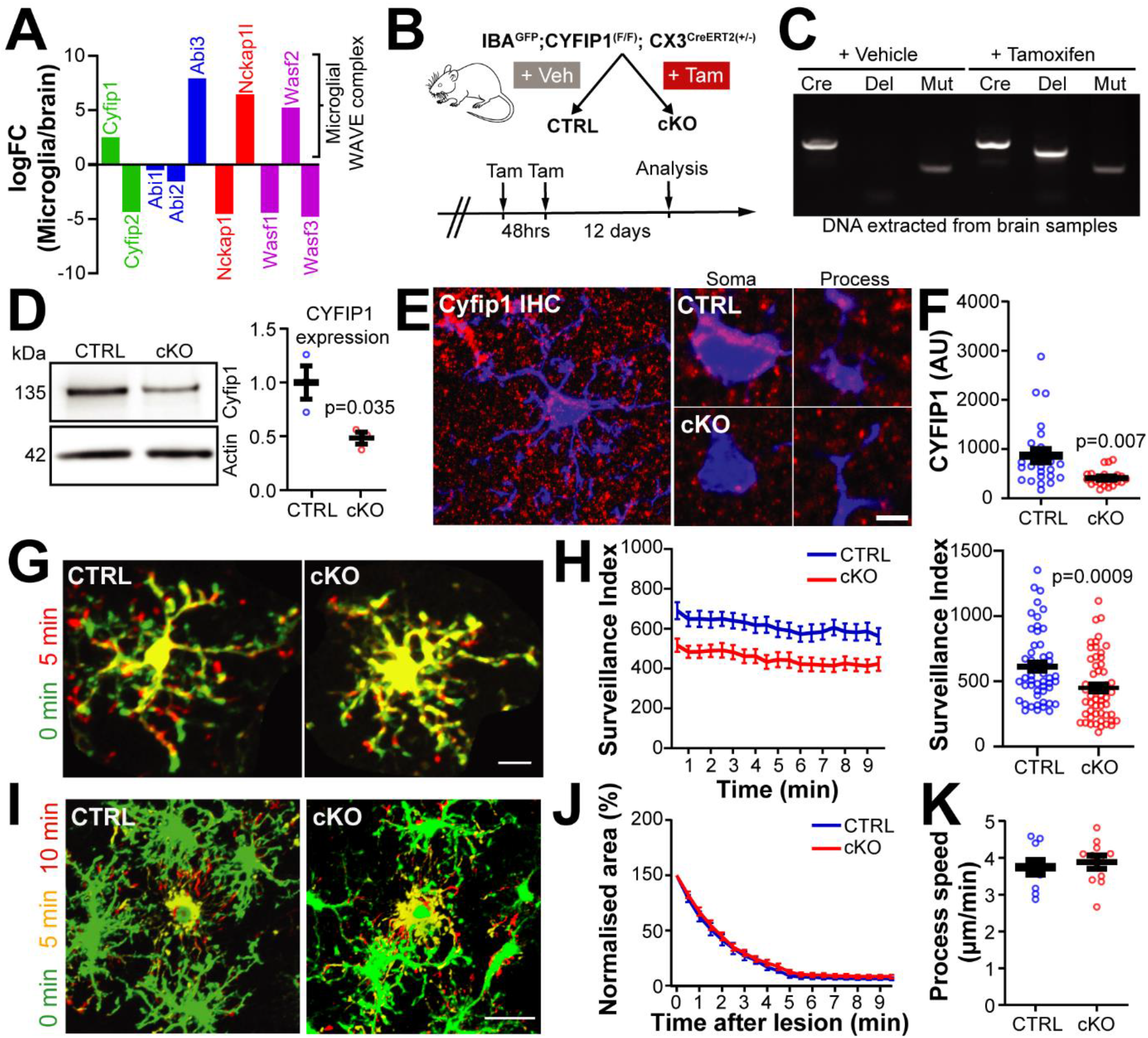
Generation of a microglia-specific Cyfip1 conditional knockout mouse model. **(A)** mRNA expression of WRC components, showing log(fold change) (logFC) of FPKM values from microglia and whole brain samples. Positive logFC values indicates enrichment in microglia. Adapted from Zhang et al. (2014). See also Fig.S3 **(B)** Schematic showing generation of conditional *Cyfip1* deletion in microglia using the Cre^ERT2^driven from the microglia-specific CX3CR1 promoter. Cre was activated through treatment of experimental mice with tamoxifen. See also Fig.S4 **(C)** PCR confirmation of recombination of the floxed *Cyfip1* cassette in control and Cyfip1 cKO mice. **(D)** Western blot for Cyfip1 in an enriched microglial extract from adult mouse brain, quantification in the right-hand panel (control: 1 ± 0.15, cKO: 0.48 ± 0.06; p<0.05, Student’s t-test, n=3 lysates preparations). **(E-F)** IHC for Cyfip1 in IBA^GFP^control and cKO mice. **(E)** Left panel: max projections of Cyfip1 staining. Right panels: zooms of soma and processes show reduction of Cyfip1 puncta in cKO microglia. Scale bar: 10 μm (large), 2 μm (zoom). **(F)** Quantification of Cyfip1 levels in the cell soma (control: 867.7 ± 132 IntDen, cKO: 410.2 ± 30.85 IntDen; p<0.05, Mann-Whitney; n=25 cells). **(G)** Superimposed timelapse images of microglia from control or cKO brain slices, showing extensions (red), retractions (green) and stable regions (yellow) over 5 minutes. Scale bar: 10 μm. **(H)** Surveillance index of cKO microglia decreased compared to controls (control: 612.5 ± 37.6 px per 30 sec, cKO: 449.8 ± 32.7 px per 30 sec; p<0.001, Mann-Whitney; n=52-56 cells). **(I-K)** Chemotaxis is normal in cKO microglia. **(I)** Superimposed timelapse showing process chemotaxis in control and cKO slices. Scale bar: 10 μm. **(J)** No change in process convergence. **(K)** Single process tracking shows normal extension speed (control: 3.77 ± 0.14 μm/min; cKO: 3.90 ± 0.10 μm/min; n= mean proceed speed values from 10-11 lesions).

To study loss of Cyfip1 in microglia *in vivo*, a conditional knockout (cKO) mouse model was generated by crossing a floxed *Cyfip1* mouse with a line expressing an inducible CreERT2 fusion protein driven by the microglia-specific *Cx3cr1* promoter (Fig. 3B, S4) (Davenport et al., 2019; Goldmann et al., 2013; Yona et al., 2013). These mice were further crossed with the IBA^GFP^ reporter line used above to visualise microglia *in situ*. Induction of *Cyfip1* recombination in this line requires delivery of tamoxifen to enable translocation of CreERT2 fusion into the nucleus. Tamoxifen was delivered orally in two consecutive 4 mg doses between P28-30 and animals were left for 14 days after initial treatment to allow for turnover of Cyfip1 protein (Fig. 3B; see Methods). Recombination of the *Cyfip1* cassette was confirmed by PCR from brain tissue from control or Cyfip1 cKO animals, and highlighted a lack of recombination in the absence of tamoxifen (Fig. 3C). To confirm Cyfip1 loss at the protein level, enriched microglia samples were extracted from control and cKO brains using a Percoll-based isolation method. Enriched microglial lysates showed a significant decrease in Cyfip1 expression of approximately 50% (Fig. 3D, p<0.05). Finally, loss of microglial Cyfip1 was shown on the single cell level by IHC of slices stained with an antibody against Cyfip1 (Fig. 3E-F). In control slices, Cyfip1 staining is punctate and clearly enriched in microglial somata but also seen within microglial processes (Fig. 3E). In contrast, the integrated density of Cyfip1 staining was significantly reduced in microglia from cKO slices (Fig.3F, p<0.01). Together, these data show that tamoxifen treatment in this model leads to specific and efficient loss of microglial Cyfip1 *in vivo*, facilitating future study.

### Cyfip1 deletion leads to impaired surveillance, morphological defects and microglial activation

We first investigated microglial motility in control and Cyfip1 cKO animals using the two assays of surveillance and chemotaxis described previously (Fig. 3G-H). Quantifying surveillance of cKO microglia compared to control showed a 24% reduction in surveillance index (Fig. 3G-H, p<0.01). Interestingly, loss of Cyfip1 had no effect on the chemotactic responses to laser-induced lesion, with control and cKO microglia responding with equal process convergence (Fig. 3I-J). In agreement with this, tracking individual processes showed no change in speed between genotypes (Fig.3K). These data show that loss of microglial Cyfip1 specifically impairs the surveillance behaviour of microglia, leaving chemotaxis unimpaired.

We hypothesised that impaired surveillance in Cyfip1 cKO microglia was due to morphological defects in these cells. To test this, control and cKO brains were fixed, and 100 μm sections were GFP signal amplified to enable 3D reconstructions of individual hippocampal microglia (Fig. 4A-C) (Peng et al., 2014). Sholl analysis revealed a decrease in the complexity of microglial processes that appeared consistent across the whole arbor (Fig. 4B). Supporting this, branching and total process length were both significantly reduced in the Cyfip1 cKO (Fig. 4C). It has recently been found that microglial processes are decorated with many highly dynamic filopodia that support surveillance of the brain parenchyma (Bernier et al., 2019). These structures were clearly visible in our fixed tissue (Fig.4D), but were not included in our initial reconstructions. To specifically investigate filopodia, a lower minimum threshold for process length (1 μm instead of 5 μm) was used for reconstructions, and the number of tips within this 1-5 μm length range was defined to represent the number of filopodia. Interestingly, the number of filopodia per cell was decreased by 38% in cKO microglia (Fig.4E, left). This effect remained even when normalised to the total process length (Fig.4E, right), suggesting that this is not simply a consequence of reduced arbor complexity seen in Fig.4C.

**Fig.4:**
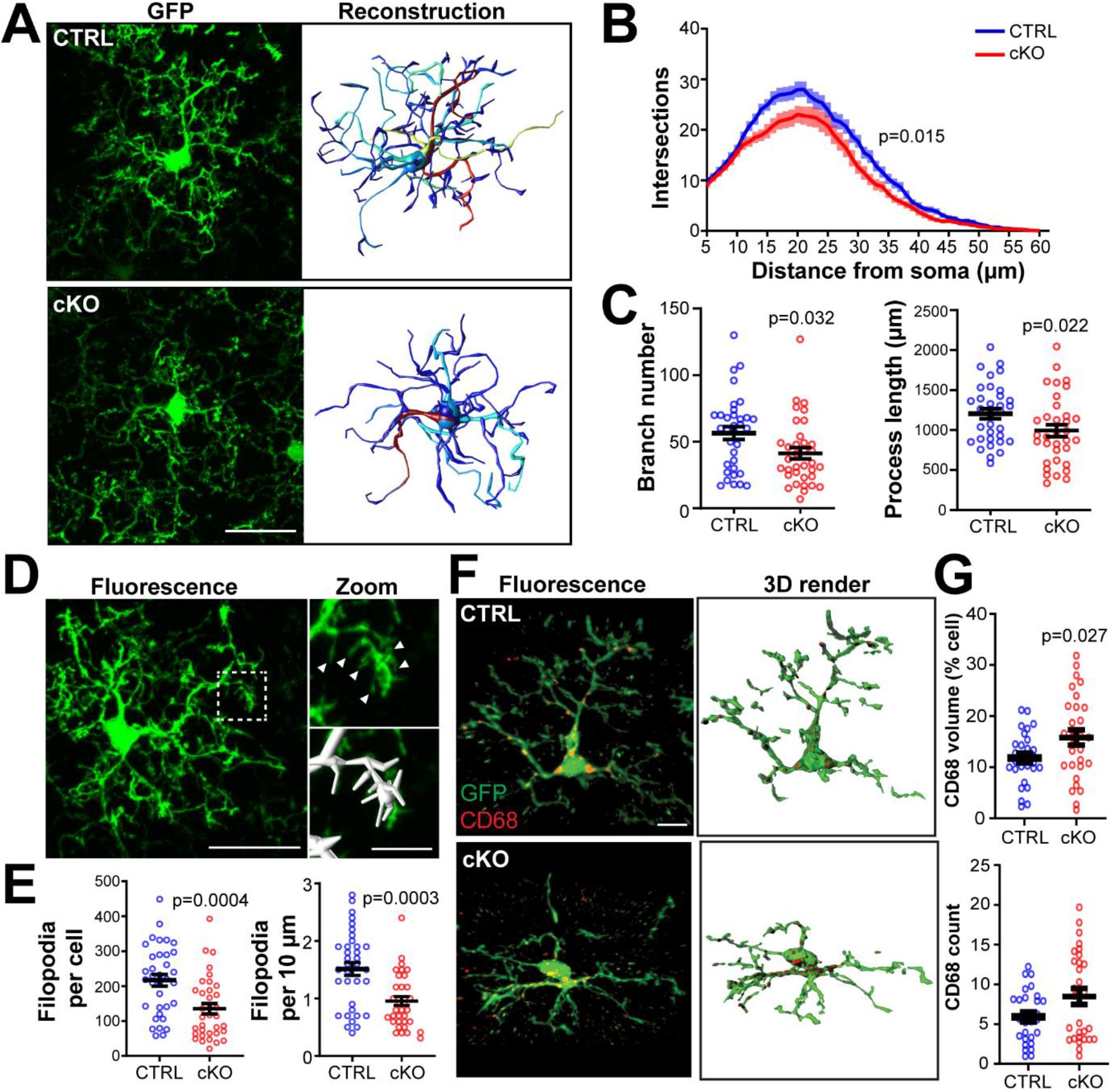
Decreased morphological complexity and altered lysosomal content of Cyfip1 cKO microglia. **(A)** Representative projections of GFP IHC (left) and 3D reconstructions (right) of microglia taken from fixed slices of control and Cyfip1 cKO brains, showing reduced complexity of arborisations. **(B)** Sholl analysis of reconstructions reveal a decrease in the number of intersections across the arbor (p<0.05, 2-way repeated measures ANOVA, n=34 cells). **(C)** Summary of morphological characteristics of microglia; reduced branch number (control: 56.44 ± 4.85, cKO: 41.31 ± 4.22; p<0.05, Mann-Whitney) and total process length (control: 1206 ± 63.30 μm, cKO: 992.6 ± 74.47 μm; p<0.05, Student’s t-test). **(D)** Microglial filopodia decorate processes (white arrows, top right) and can be automatically reconstructed by Vaa3D (bottom right). **(E)** Quantification of the number of filopodia shows reductions in cKO cells both when measured per cell (control: 216.9 ± 16.78, cKO: 135.1 ± 14.96; p<0.001, Mann-Whitney) or per 10 μm of process (control: 1.51 ± 0.11, cKO: 0.96 ± 0.08; p<0.001, Mann-Whitney). n=35 cells. Filopodia we defined as process tips with length <5 μm. **(F-G)** CD68 staining in cKO microglia. **(F)** Example images of immunofluorescence (left) and 3D volume renders (right) of CD68 staining in single microglia. **(G)** Quantification reveals increase in CD68 burden in cKO cells as a percentage of total cell volume (control: 11.79 ± 0.96 %, cKO: 15.82 ± 1.48 %; p<0.05, Student’s t-test, n=28-30 cells).

Finally, we wanted to investigate whether loss of Cyfip1 had any broader impact on microglial function. The activation state of microglia alters many aspects of their physiology, including phagocytosis, cytokine production and gene expression (Bennett et al., 2016; Wolf et al., 2017). The scavenger receptor CD68 (also known as macrosialin), a monocyte-specific marker of the lysosome, is commonly used as a readout of microglial activation (Hong et al., 2016; Lui et al., 2016; Qin et al., 2018) (Fig.4F). To see if loss of Cyfip1 alters lysosome numbers in microglia, hippocampal sections were stained for CD68 (lysosomes) and GFP (cell fill) and the lysosomal content of microglia was quantified, based on a published protocol (Schafer et al., 2014). Interestingly, Cyfip1 cKO microglia had an increased volume of CD68 positive structures as a percentage of the total cell volume (Fig.4G, p<0.05). This effect appeared to be mostly driven by an increased number of lysosomes per cell. This suggests that loss of Cyfip1 increases the activation state of microglia. Together with the morphology and surveillance defects described, these data highlight the importance of Cyfip1 for normal microglial physiology and relevance to disease.

## Discussion

Control of microglial shape and motility is critical to facilitate these cells’ multitude of interactions during health and disease. Here, we report that microglial ramification and process morphology is dependent on sustained activity of the actin branching complex Arp2/3. Inhibiting Arp2/3 led to a specific defect in surveillance of the brain parenchyma, without affecting chemotaxis. Moreover we demonstrate that the schizophrenia and autism-associated *Cyfip1*, a core component of the WRC and a key upstream regulator of Arp2/3, also controls this pathway controlling microglial morphology and surveillance. We demonstrate that disrupting actin remodeling by Cyfip1 leads to broader changes in microglial activation.

The ramified shape of surveying microglia forms dynamically from the extension and retraction of processes (Arcuri et al., 2017; Madry and Attwell, 2015). Furthermore, in response to chemotactic signals such as ATP/ADP release from damaged cells, microglia rapidly alter their morphology, extending nearby processes towards the signal. Cytoskeletal dynamics are tightly controlled by a network of actin-associated proteins the functions of which in microglia remain largely unknown. As mutations in many of these genes are linked to neurological conditions, it is important to understand how actin regulators control microglial behavior and contribute to disease. By analysing published gene expression data, we identified an enrichment in microglia of components of the WAVE regulatory complex (Wasf2, Cyfip1, Nap1l, Abi3) that activates Arp2/3 to form new branched actin filaments (Galatro et al., 2017). The importance of Arp2/3 activity in microglia is highlighted by the dramatic changes in cell morphology observed in brain slices treated with CK666 to block this complex. Reduced microglial ramification and dysmorphic process blebbing was observed after only 30 minutes of drug exposure, implying that Arp2/3-mediated actin branching is tonically active in surveying microglia. Higher-order processes are mainly responsible for surveillance over the timeframe that we imaged for and were preferentially affected by Arp2/3 inhibition, leading to severely impaired surveillance of the brain parenchyma. The bleb-like processes seen during Arp2/3 and Rac1 inhibition are reminiscent of alternative actin-based protrusions used by ameboid cells to migrate through 3D environments. Indeed, blebbing is stimulated by inhibition of Arp2/3 and dependent on RhoA/ROCK activity in other cells, suggesting that the dynamics of microglial actin are set by a balance of different RhoGTPase-dependent pathways (Paluch and Raz, 2013; Rotty et al., 2017; Xu et al., 2016).

Cyfip1 has well-established roles in Arp2/3 complex activation, via its presence in the WRC (Chen et al., 2010; Krause and Gautreau, 2014). Here we used a novel conditional knockout model to show that deletion of Cyfip1 leads to impaired surveillance of the brain parenchyma and reduced morphological complexity of microglia, strongly supporting a role for Cyfip1 in controlling their resting state behaviour. As these phenotypes are similar to those that we report during inhibition of the actin branching machinery, we hypothesise that Cyfip1 deletion leads to disruption of the WAVE regulatory complex and Arp2/3 activation. In neurons, Cyfip1 also functions as a repressor of cap-dependent translation in a complex with the Fragile X mental retardation protein (FMRP) (Napoli et al., 2008). Whilst we cannot exclude a role for altered protein translation in our Cyfip1 cKO model, FMRP is not expressed in adult hippocampal microglia, making this unlikely (Gholizadeh et al., 2015). We also report reduced filopodia numbers in cKO cells, even after accounting for altered ramification of the cell’s major processes. A cAMP-dependent mechanism governing filopodia formation in microglia was recently discovered (Bernier et al., 2019). Whether Cyfip1 functions downstream of cAMP signaling to control filopodia formation is currently unknown, although the major cAMP effector, PKA, interacts with Rac1 and Wave2 to control membrane protrusion (Howe, 2004; Yamashita et al., 2011).

Both Arp2/3-inhibition and Cyfip1 cKO microglia specifically impact surveillance over chemotaxis, possibly because the processes extending to the lesion in chemotaxis are already oriented towards the lesion site and thus require only filament elongation and not branching. Our observations fit with other studies showing divergence in the signaling pathways underlying surveillance and chemotaxis; the potassium channel THIK-1 controls surveillance but is dispensable for chemotaxis, conversely the ADP receptor P2Y_12_ is only required for chemotaxis (Bernier et al., 2019; Haynes et al., 2006; Madry et al., 2018b; Sipe et al., 2016). Given that both forms of motility remains critically dependent on actin, as seen in our cytochalasin D treatment, it is remarkable that the divergence of signalling underlying these behaviours extends down to the machinery regulating actin filament formation.

A key advantage of our cKO model is the ability to discern microglial-specific effects of Cyfip1 deletion. Most *in vivo* studies of Cyfip1 have relied on heterozygous models, where Cyfip1 expression is altered in all cells, making inferring cell autonomy of phenotypes extremely challenging (Bozdagi et al., 2012; Pathania et al., 2014; De Rubeis et al., 2013). This is highlighted by two recent studies on rodent Cyfip1 haploinsufficient models that both reported myelin thinning, but drew different conclusions as to whether altered neuronal activity or oligodendrocyte function underpinned these effects (Domínguez-Iturza et al., 2019; Silva et al., 2019). Indeed, microglia are in active and intimate association with neurons during key stages in development, which helps to shape neuronal connectivity (Miyamoto et al., 2016; Schafer et al., 2012; Squarzoni et al., 2014; Wake et al., 2013). These interactions occur within the surveillance field of a microglial cell and could well be affected by alterations in the cell’s ramification and motility. Thus, discerning whether altered microglial function in Cyfip1 models affects neuronal growth, connectivity or behavioral phenotypes is a key question arising from our work.

Interestingly, other WRC components expressed in microglia are increasingly implicated in neurological disorders with microglial pathologies. WASF2 was recently found to be differentially expressed in microglia from ASD individuals (Velmeshev et al., 2019). ABI3, which interacts directly with CYFIP1, is exclusively expressed by microglia in the brain and is a risk gene for Alzheimer’s disease (Satoh et al., 2017; Sims et al., 2017). Similarly LRRK2 is a Parkinson’s disease (PD)-associated gene that regulates WAVE2-dependent phagocytosis of dopaminergic neurons by microglia in PD models (Kim et al., 2018). Thus, we provide direct support for a growing body of evidence suggesting that disruption of the WAVE regulatory complex is implicated in neurological conditions, and illustrate the need to investigate the cell autonomous roles of these genes to properly understand their contribution to pathology.

Aberrant activation of microglia has been strongly implicated in autism and schizophrenia (Bilbo et al., 2018; Bloomfield et al., 2016; Deakin et al., 2018; Sellgren et al., 2019). We find loss of microglial Cyfip1 increased CD68 positive content, a marker of microglial activation, providing a link between the clinical associations and microglial functions of Cyfip1 (Lui et al., 2016; Qin et al., 2018). Loss of microglial ramification and reduced surveillance are characteristic features of activated microglia, though the factors controlling cytoskeletal remodeling during activation are largely unknown (Hines et al., 2013; Paris et al., 2018). Our data suggest that reduced Cyfip1 and Arp2/3 signalling could contribute to altered morphology during activation. Interestingly, downregulation of Arp complex-related genes has been reported in several contexts where microglia are activated; microglial Cyfip1 levels are reduced in LPS-infused mouse brain, and RNA sequencing from both human and mouse microglia in ageing cortex showed a downregulation of Arp2/3 components ARPC1A and ARPC1B, alongside CYFIP1 and WASF2 (Galatro et al., 2017; Pan et al., 2020; Zhang et al., 2014). Whether altered expression of these genes augments microglial activation in pathological contexts remains to be seen. Overall, this study adds to our growing understanding of the fundamental biology underpinning microglial morphology and surveillance, and identifies novel implications for neuropsychiatric disease.

## Acknowledgements

Supported by grants to J.T.K from the UK Medical Research Council (G0802377, MR/N025644/1) and ERC (282430). An MRC funded 4-year Clinical Neurosciences PhD programme to J.D, Marie Curie Fellowship to R.J and Wellcome Trust Senior Investigator Awards 099222 and 219366 to D.A. We thank Stuart Martin for genotyping, and Christian Coville-Cooke and Davor Ivankovic for advice and comments on the manuscript.

The authors declare no conflict of interest.

## Author contributions

J.T.K. and J.D. conceived the paper. J.D. carried out all experiments and analysis. I.L.A.-C. assisted in developing imaging methods and drug perfusion system, and developed the 3D Sholl analysis software. R.J. developed the surveillance and chemotaxis analysis software. J.T.K. and G.L-D. generated the Cyfip1 cKO mouse line. D.A. provided input to microglial imaging experiments and analysis. J.T.K. and J.D. wrote the manuscript.

## Software

Code for analysing microglial morphology is freely available at https://github.com/AttwellLab

## Supplementary figures

**Fig.S1:**
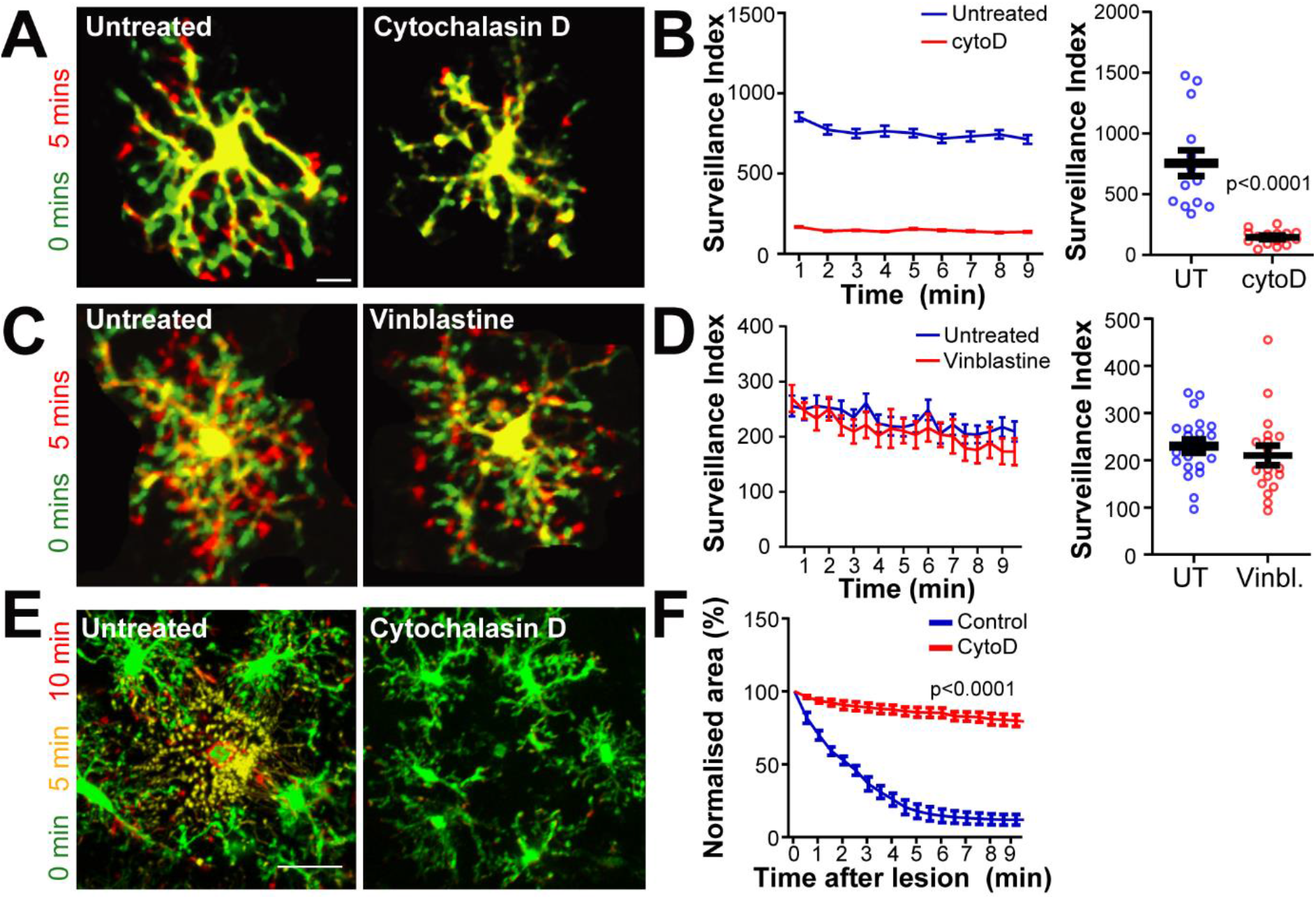
Microglial motility requires actin but not microtubule polymerisation. **(A-B)** 10 μM cytochalasin D treatment abolishes microglial surveillance **(A)** Superimposed maximum intensity projection of representative cells at 0 and 5 minute timepoints (green = retraction; red = extension; yellow = stable). **(B)** Average of surveillance index over the course of each movie (untreated: 755 ± 106 px per 30 sec, cytochalasin D: 145 ± 15 px per 30 sec, p<0.0001, Student’s t-test, n=14-15 cells). **(C-D)** Depolymerisation of microtubule cytoskeleton with 10 μM vinblastine does not affect microglial surveillance (untreated: 229 ± 14 px per 30 sec, vinblastine: 210 ± 21 px per 30 sec, n=18-21 cells). **(E)** Binarised max projections of untreated and cytochalasin D microglia. Convergence of processes seen as a reduction of red polygon area that is unchanged in cytochalasin D condition. **(F)** Change in area (normalised to starting area) of polygon over time (p<0.0001, 2-way repeated measures ANOVA, n=7 lesions).

**Fig.S2:**
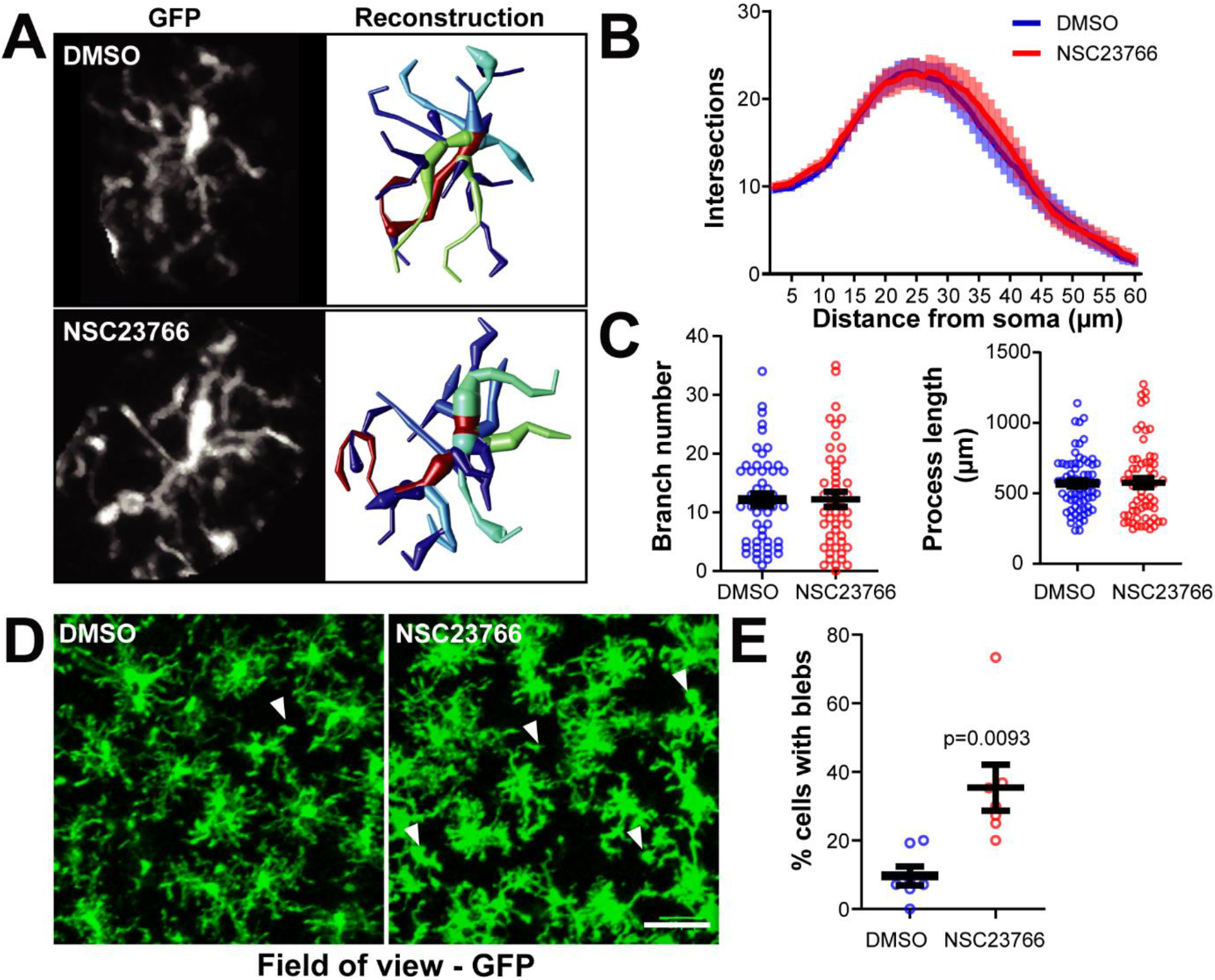
Inhibition of Rac1 increases process blebbing. **(A)** Representative examples of GFP signal (left) and 3D reconstructions (right) of single microglia from DMSO and NSC23766 treated slices. **(B)** Sholl analysis of intersections from 3D reconstructions. **(C)** No change in branch number or total process length between conditions. n=46-48 cells per condition. **(D)** Field of view of DMSO and NC237666 treated slices showing presence of blebs in drug-treated condition (right, white arrows). Scale bar: 25 μm. **(E)** Quantification of proportion of microglia with blebs (DMSO: 9.7 ± 2.8%, NSC23766: 35.4 ± 6.7%, p<0.01, Student’s t-test, n=7 fields of view).

**Fig.S3:**
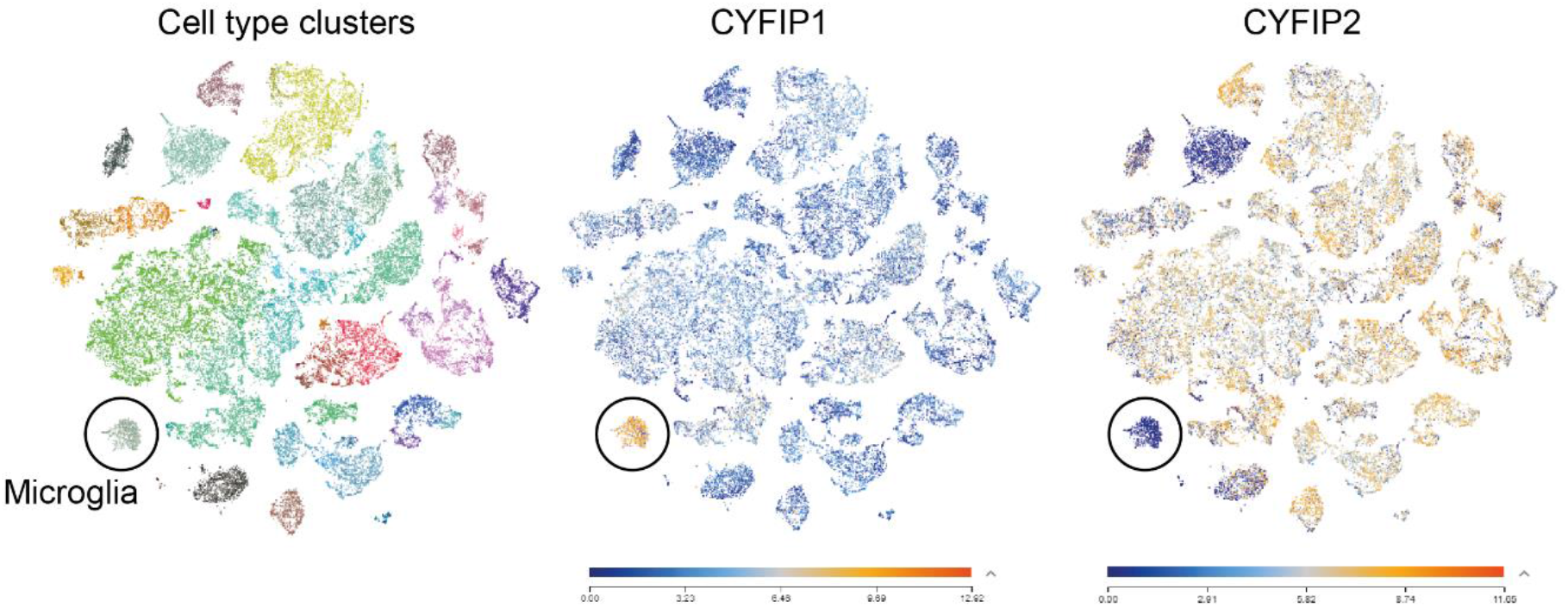
CYFIP1 is the predominant CYFIP protein in human microglia. t-SNE plots of single cell sequencing data from human cortex from the Allan Brain Map by the Allan Institute (https://celltypes.brain-map.org/rnaseq/human/cortex, accessed 10.04.2020). **(A)** Colour mapping highlights clusters identifying specific cell populations, including clearly distinct microglia cluster (circle). **(B-C)** *CYFIP* gene expression mapped onto cell clusters, showing that microglia are specifically enriched in *CYFIP1* **(B)** over *CYFIP2* **(C)**. Data from: Hodge, R.D., Bakken, T.E., et al. (2019).

**Fig.S4:**
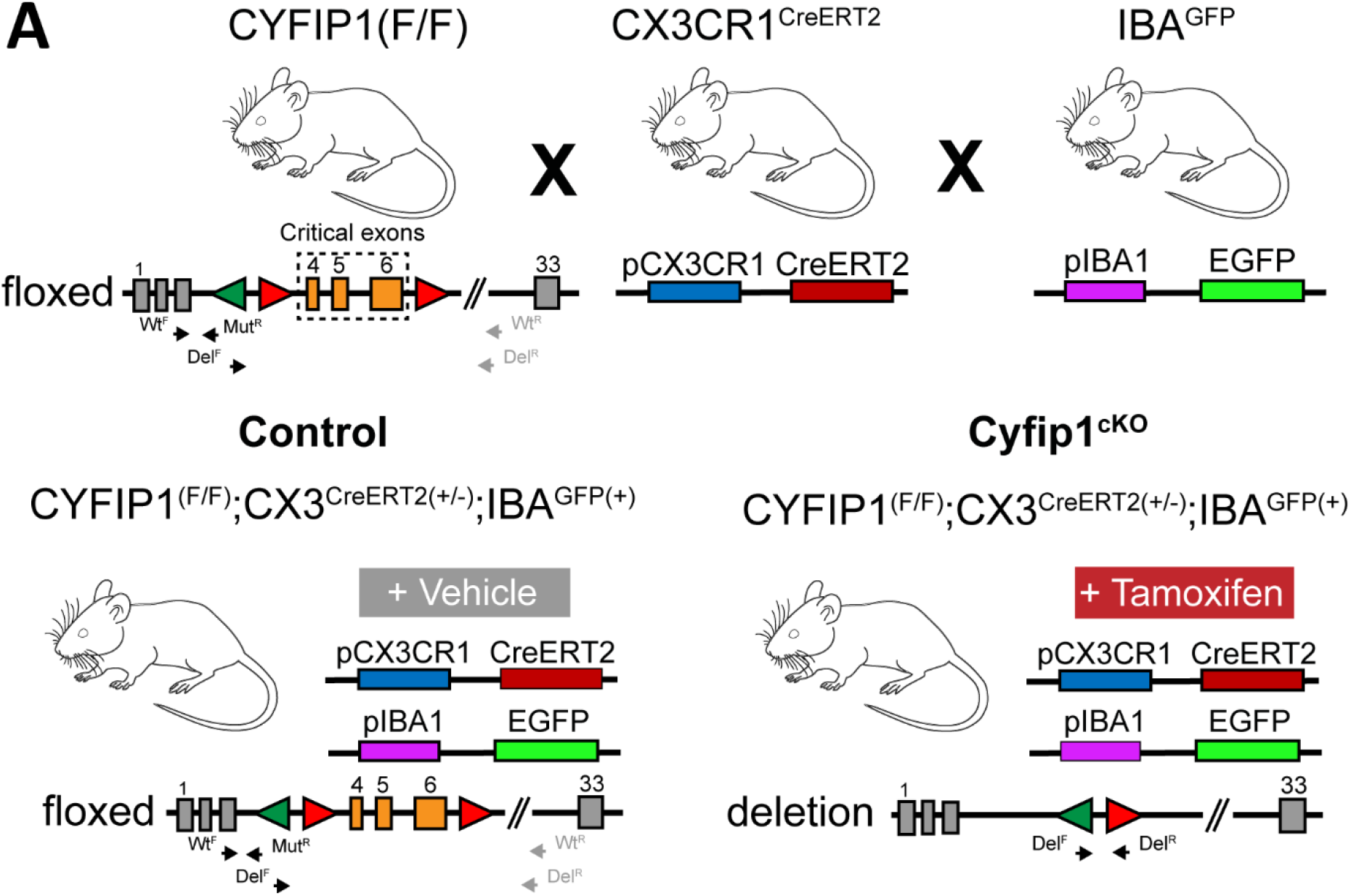
Generation of Cyfip1 conditional knockout mouse model. **(A)** Schematic illustrating the genetic modifications used to generate the *Cyfip1* conditional knockout (cKO) mouse model. A floxed *Cyfip1* line was generated using a knockout-first strategy and bred to homozygosity in all parental mice. To generate knockout alleles, floxed *Cyfip1* mice were is crossed a knock-in CX3Cr1^CreERT2^ line. CX3Cr1^CreERT2^ allele was kept heterozygous in all breeding and experimental animals. An IBA^GFP^line used previously was crossed in to fluorescently label microglia. Recombination of the floxed *Cyfip1* locus excises critical exons 4-6, leading to loss of functional *Cyfip1*. Nuclear translocation of Cre recombinase was activated by tamoxifen treatment and was inactive in vehicle treatment. Small arrowheads identify approximate sites for PCR primer sequences for wildtype (Wt^F^/Wt^R^), conditional (Wt^F^/Mut^R^) and the deletion (Del^F^/Del^R^) alleles of the *Cyfip1* locus. Grey arrows denote no PCR product.

## Materials and methods

### Key resources table

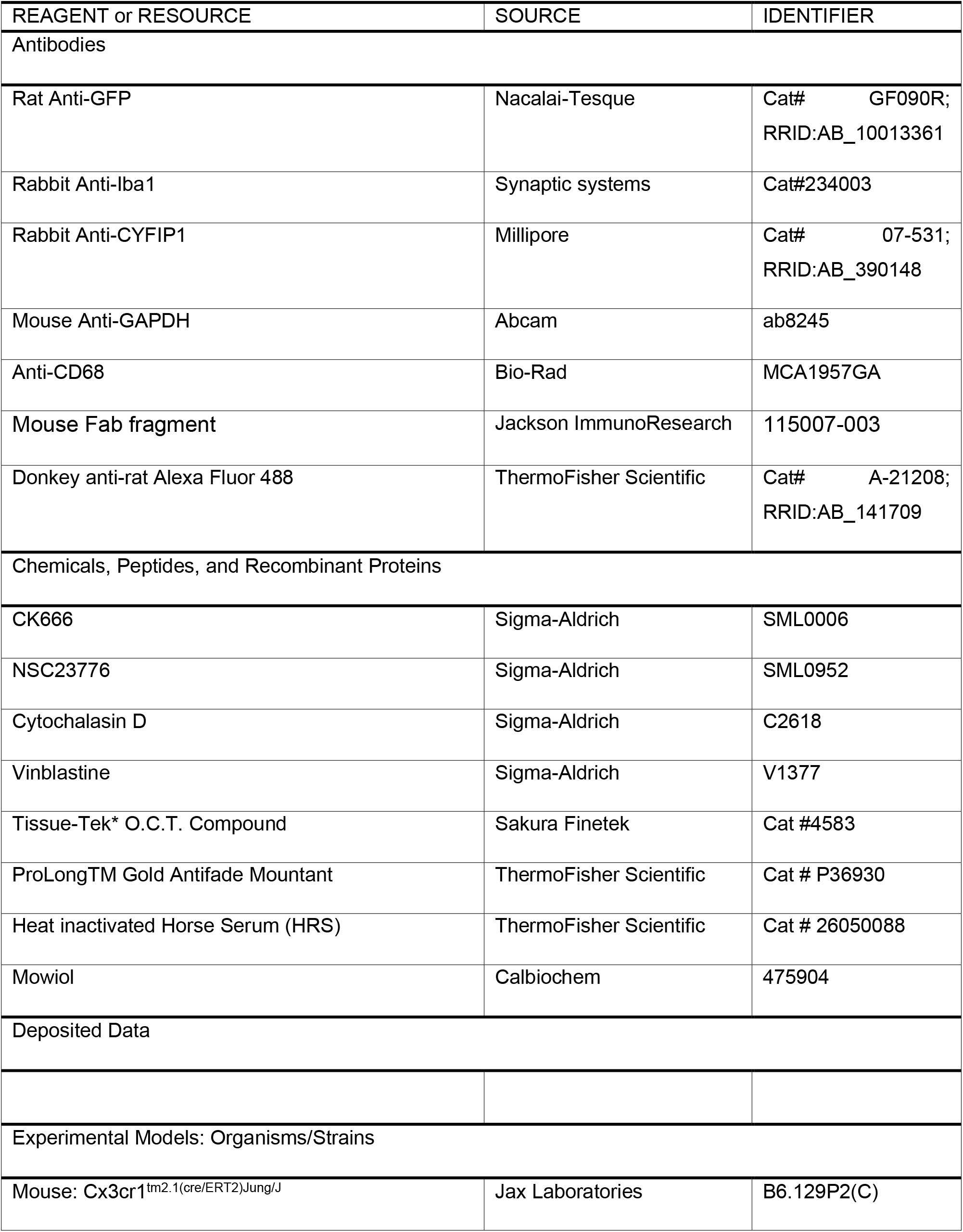
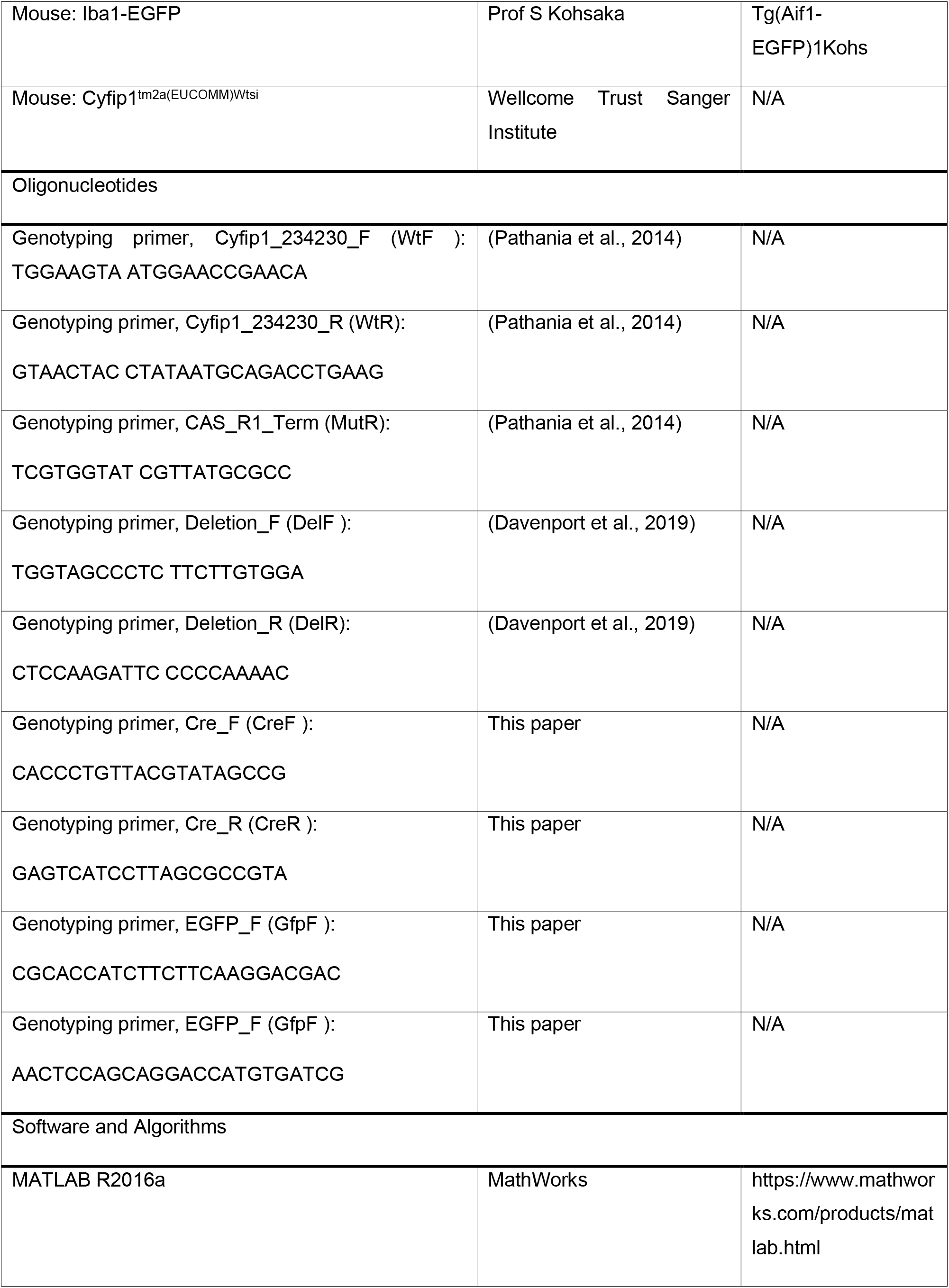
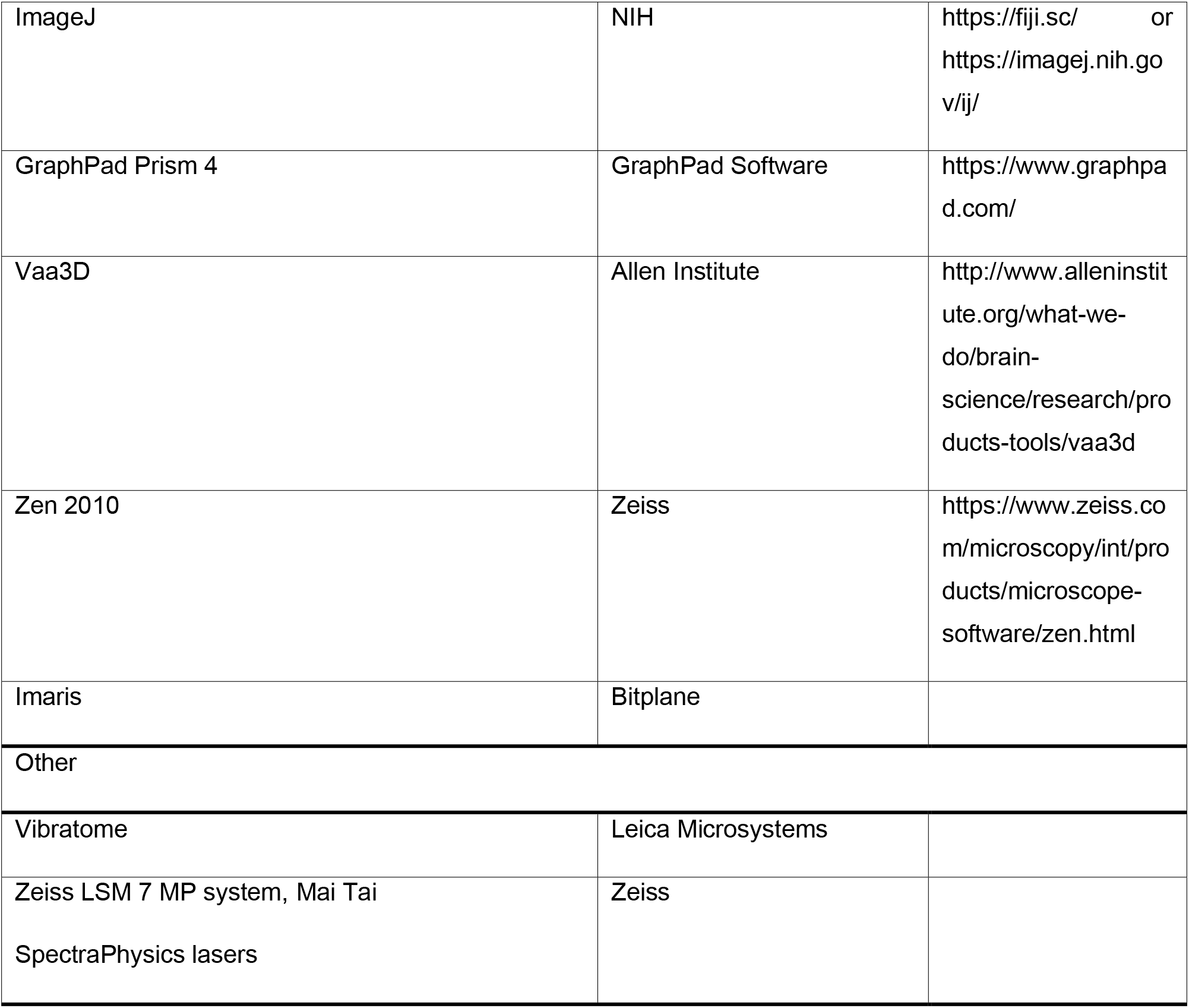

#### Contact for reagent and resource sharing

Further information and request for resources and reagents should be directed to and will be fulfilled by the Lead Contact, Josef Kittler.

#### Experimental model and subject details

##### Animals

Animals of either sex were used for all experiments. Ages are stated in the figure legends for each experiment. All procedures for the care and treatment of animals were in accordance with the Animals (Scientific Procedures) Act 1986 and had full Home Office ethical approval. All animals were maintained under controlled conditions (temperature 20 ± 2°C; 12 h light-dark cycle). Food and water were provided ad libitum. Animals were group housed in conventional cages and had not been subject to previous procedures.

##### Mouse line generation and inducible Cre recombinase activation

The *Cyfip1* knockout (KO) mouse line (MDCK; EPD0555_2_B11; Allele: Cyfip1^tm2a(EUCOMM)Wtsi^) was generated using the Knockout-First strategy on C57BL/6N Taconic USA background and was obtained from the Wellcome Trust Sanger Institute as part of the International Knockout Mouse Consortium (IKMC) (Skarnes et al., 2011). The KO-first allele contains an L1L2_Bact_P cassette flanked by frt sites inserted 50 of critical exons 4 to 6 of *Cyfip1*, disrupting gene function. The cassette can be deleted in the presence of flp recombinase by crossing with a flp-expressing strain. The flp-directed recombination produces a functional allele with the critical exons flanked by loxP sites (*Cyfip1^F^*^/F^). *Cyfip1* conditional knockout animals were generated by crossing *Cyfip1^F^*^/F^ animals with the Cx3cr1^tm2.1(cre/ERT2)^ line. Microglia were visualised by crossing conditional knockout animals with an Iba1^GFP^ line. Experimental mice were generated as a result of *Cyfip1^F^*^/F^ x *Cyfip1^F^*^/F:Cx3cr1-Cre(+/-):IBA-GFP^ crosses to produce littermate controls. All surveillance imaging was done using Iba1^GFP^ labeled mice.

To activate Cre in the Cx3cr1^(Cre/ERT2)^line, estrogen receptor agonist tamoxifen was administered to adult mice (P28-P40) via oral gavage. Tamoxifen was dissolved in 40 mg/mL solution of 90% cornoil (Kolliphor EL, Sigma):10% ethanol by sonication and two doses of 4 mg/100 μl were administered, 48 hours apart. Tamixfen solution was made fresh for each dosing regimen and kept at 4°C between doses.

##### Acute slice preparation

To prepare acute hippocampal slices, male and female mice aged postnatal day 28-34 were used. Immediately after decapitation, the brain was removed and kept in ice-cold dissecting solution (in mM: 87 NaCl, 25 NaHCO_3_, 10 glucose, 75 sucrose, 2.5 KCl, 1.25 NaH_2_PO_4_, 0.5 CaCl_2_ and 7 MgCl_2_) saturated with 95%O_2_/5% CO_2_. Transverse hippocampal slices (300 μm) were obtained using a vibratome (Leica, VT–1200S). Slices were transferred to HEPES-buffered ACSF (in mM: 140 NaCl, 10 glucose, 10 HEPES, 2.5 KCl, 1 MgCl_2_, 2 CaCl_2_, 1 NaH_2_PO_4_, saturated with 100% O2 (pH 7.4) at 22◦C, and left to recover for 30 minutes prior to use.

##### Isolation of enriched microglia fraction

To generate protein lysates of enriched adult microglia from transgenic mice, a Percoll-based isolation method was used as described previously (Lee and Tansey 2013). Briefly, 3 mice per experimental condition were transcardially perfused with ice-cold PBS, brains extracted and finely minced using a flat scalpel blade in 3 ml HBSS. Tissue was enzymatically digested in a dispase-papain-DNAase solution and triturated into a single cell suspension. The suspension was then loaded into a 70%:37%:30% Percoll/HBSS gradient. After centrifugation without brakes, myelin debris migrated to the 30%:37% interface and microglial cells were enriched in the 70%:37% interface. The microglia fraction was extracted, washed and lysed in sample buffer.

### Methods details

#### Transcardial perfusion fixation and immunohistochemistry

Brains were fixed by transcardial perfusion (TCP) of 4% paraformaldehyde (PFA) (4% PFA, PBS). Animals underwent terminal anaesthesia via intraperitoneal injection of 1 μl/g pentobarbital solution. When fully anaesthetised, 10-15 ml of ice-cold 4% PFA was perfused into the left ventricle using a peristaltic pump, the animal decapitated, brain removed and placed in ice-cold 4% PFA overnight. Certain protocols required TCP of PBS to remove all blood from the brain. In these cases, the protocol above was used except that ice-cold PBS was perfused.

Microglial morphology in *Cyfip1* cKO animals was assessed using perfusion-fixed brains from litter matched pairs. Brains were transferred to PBS and 100 μm slices made using a vibratome (Leica Microsystems, Heerbrugg, Switzerland). For immunohistochemistry, free floating sections were washed in PBS before permeabilization in blocking solution (10% horse serum, 0.5% BSA, 0.2% Triton X-100, PBS) for 2-4 hours and incubation with primary antibody diluted in block solution overnight at 4◦C. For mouse primary antibodies, slices were first incubated overnight at 4°C with mouse Fab fragment (1:50 with blocking solution). Slices were washed 4-5X in PBST for 2 hours then incubated for 3-4 hours with secondary antibody at RT. Slices were then washed 4-5× in PBS for 2 hours and mounted onto glass slides using Mowiol mounting medium. For cryoslices, brains were dehydrated with 30% sucrose/PBS solution and frozen at −80°C before embedding in OCT (Optimal Control Temperature). 30 μm sections made using a Cryostat (Bright Instruments, Luton, UK) and stored at −20°C as free-floating slices in cryoprotect solution (30% ethylene glycol, 30% glycerol, 40% PBS). For antigen retrieval, slices were incubated in sodium citrate solution at 80°C for 40 mins and then washed 3× in PBS prior to blocking.

#### Quantifications of microglial morphology

Microglial morphology in *Cyfip1* cKO animals was assessed using perfusion-fixed brains from litter matched pairs. Thick sections underwent IHC (see Transcardial perfusion fixation and immunohistochemistry) to amplify GFP signal. High resolution z-stacks of single microglia were taken using a Zeiss LSM700 upright confocal microscope using a 63X oil objective (NA: 1.4) (70-100 μm z-stacks; xy resolution: 0.10 μm; z step: 0.32 μm, 1024×1024 px). Input z-stack images were pre-processed using a custom ImageJ macro to subtract background (50 px), median filter (2 px), and improve contrast (manual adjustment). Semi-automated 3D reconstructions were produced using the ‘APP2’ tracing plugin in Vaa3D software (http://home.penglab.com/proj/vaa3d), and manually checked with any clear abnormalities corrected. Morphological parameters of reconstructions were extracted using custom MATLAB code (available at https://github.com/AttwellLab). For morphology data in drug treatment experiments, the first frame of each live imaging movie was used as the raw data for generating reconstructions described above.

#### Two-photon imaging and drug treatments

Acute slices (see Acute slice preparation) of IBA^GFP^ animals were imaged in continually perfused HEPES-buffered ACSF using a two-photon microscopy system (Zeiss LSM 7 MP system, Mai Tai SpectraPhysics lasers). Slices were imaged between 30 minutes to 4 hours after slicing. Hippocampus was found using brightfield illumination and regions containing well-labelled, unactivated microglia between 50-150 μm from the slice surface were used for timelapse recordings. For surveillance measurements, z-stacks of 52 μm regions (26 stacks, 2 μm interval, 512×512 px) were taken every 30 s for 10 minutes, except for cytochalasin D treatment which was taken every 60 s for 10 minutes. CK666 and NSC23766 drug treatments were imaged at 1.5× zoom and *Cyfip1* cKO at 2X zoom. For chemotaxis measurements, a circular ROI of 15 μm diameter was selected in an area surrounding by labelled microglia. High-powered laser excitation (90% power) within the ROI was used to create a lesion in the slice, using the bleaching plugin in Zen 2010 (12 iterations, 2 speed). Immediately after lesion, 3D movies were taken using the same imaging parameters as for surveillance (2X zoom).

For cytochalasin D, slices were incubated for 10 minutes in 10 μM cytochalasin D prior to imaging in normal ASCF. For DMSO (1%), CK666 (200 μM) and NSC23776 (100 μM) treatments, acute slices were incubated for 30 minutes prior to imaging in a low-volume incubation chamber with drug solutions in HEPES-buffered ACSF. Slices were then imaged for a maximum of 30 minutes using a re-perfusion system that enabled recycling of drug-containing imaging solution.

### Quantification and statistical analysis

#### Analysis of live imaging data

For chemotaxis analysis, maximum intensity projections (MIPs) of raw images were pre-processed using a custom ImageJ macro that involved: XY registration, background subtraction (50 px), maximum filtering (2 px). As part of z-registration, single z-planes at the extremes of the stack were sometimes removed. Pre-processed MIPs were manually thresholded and the resulting movies processed using a custom previously published MATLAB script (Madry et al., 2018). Briefly, after the user manually selected the lesion region, the algorithm divides the surrounding area into concentric circles with radii at 2 μm intervals, and then segments these circles into 32 radial sectors, thus creating 32 patches between every two consecutive concentric circles. Then, for each frame, starting from the center, the algorithm searches in every radial sector for the first patch containing > 10 positive pixels (labeled microglia). The outputs of the algorithm are, for each frame, (i) the distance to the microglial process front in each sector and (ii) the surface area contained within the converging microglial process front.

For surveillance analysis, regions containing single cells were cropped from raw images. Crops were pre-processed using a separate ImageJ macro that involved: XYZ registration, background substraction (50 px), maximum filtering (2 px) and 3D-bleach correction. Finally, GFP signal from surrounding cells was removed using the Clear function in ImageJ to isolate motility of a single cell. MIPs from these processed images were binary thresholded and processed in a MATLAB script published in Madry et al. 2018. For each movie, starting with the second frame, we subtracted from each binarised frame F_t_ the preceding frame F_t-1_ and created two binarised movies, PE consisting of only the pixels containing process extensions (F_t_-F_t-1_ > 0) and PR consisting of only the pixels containing process retractions (Ft-F_t-1_ < 0). In both PE and PR, all other pixels are set to 0. The surveillance index at each timepoint was defined as the sum of PE and PR, and an average of surveillance index across the movie used to generate a single value for each cell.

#### Microglial lysosome content

Analysis of CD68 positive lysosomes in individual microglia was performed based on a published protocol by Schafer et al. (Schafer et al., 2014). Briefly, thick brain sections were antibody stained for CD68 and GFP and individual hippocampal microglia were confocal imaged with the same parameters used for microglial morphology analysis (70-100 μm z-stacks; xy resolution: 0.10 μm; z step: 0.32 μm, 1024×1024 px). Images were preprocessed in ImageJ to remove background before CD68 and GFP signal was used to surface render lysosome and microglia cell volumes respectively, using Imaris software. Particle analysis of CD68 volume render was used to calculate total volume and counts of lysosomal structures, and was normalised to total volume of the cell render were required.

#### Statistical Analysis

Volcano plot in Fig.1A using the WebGestalt online tool (http://www.webgestalt.org/) together with data from Supp. Table 1. All data were obtained using cells from at least three independent preparations or litter-matched experimental pairs. Repeats for experiments are given in the figure legends as N numbers and refer to number of cells unless otherwise stated. All statistical analysis was carried out using GraphPad Prism (GraphPad Software, CA, USA). Data was tested for normal distribution to determine the use of parametric (student’s unpaired t-test, 2-way ANOVA) or non-parametric (Mann-Whitney) tests. For posthoc analyses of two-way ANOVA, Bonferroni posthoc tests were carried out unless stated otherwise in figure legends. Data are shown as mean ± standard error of the mean (SEM).

